# The long non-coding RNA *lnc-HLX-2-7* is oncogenic in group 3 medulloblastomas

**DOI:** 10.1101/2020.06.08.140251

**Authors:** Keisuke Katsushima, Bongyong Lee, Haritha Kunhiraman, Cuncong Zhong, Rabi Murath, Jun Ying, Ben Liu, Alexandra Garancher, Ignacio Gonzalez-Gomez, Hector L. Monforte, Stacie Stapleton, Rajeev Vibhakar, Chetan Bettegowda, Robert J. Wechsler-Reya, George Jallo, Eric Raabe, Charles G. Eberhart, Ranjan J. Perera

**Author notes:** **Correspondence to:** Ranjan J. Perera, PhD, Department of Oncology, Sidney Kimmel Comprehensive Cancer Center, School of Medicine, Johns Hopkins University, 1650 Orleans St., Baltimore, MD 21231. **Authorship statement:** K.K, B.L, and R.J.P designed the study. K.K, B.L, and H.K, performed the experimental work. C.Z., B.L., R.M., and J.Y. performed data analyses. C.G.E., E.R., C.B., S.S., G.J., A.G., I.G., H.L.M., R.V., and R.W. provided cell lines, patient samples, TMAs, FFPE sections, and PDXs for the study. K.K., R.J.P., E.R., and C.G.E. wrote the main draft of the text. All authors revised and approved the final version of the manuscript.

## Abstract

**Background:** Medulloblastoma (MB) is an aggressive brain tumor that predominantly affects children. Recent high-throughput sequencing studies suggest that the non-coding RNA genome, in particular long non-coding RNAs (lncRNAs), contributes to MB sub-grouping. Here we report the identification of a novel lncRNA, *lnc-HLX-2-7*, as a potential molecular marker and therapeutic target in group 3 MBs.

**Methods:** Publicly available RNA sequencing (RNA-seq) data from 175 MB patients were interrogated to identify lncRNAs that differentiate between MB subgroups. After characterizing a subset of differentially expressed lncRNAs *in vitro* and *in vivo*, the group 3-enriched lncRNA *lnc-HLX2-7* was deleted by CRISPR/Cas9 in the MB cell line D425 Med. Intracranially injected tumors were further characterized by bulk and single-cell RNA-sequencing.

**Results:** *lnc-HLX-2-7* is highly upregulated in group 3 MB cell lines, patient-derived xenografts, and primary MBs compared to other MB sub-groups as assessed by qRT-PCR, RNA-seq, and RNA fluorescence *in situ* hybridization (FISH). Depletion of *lnc-HLX-2-7* with antisense oligonucleotides or CRISPR/Cas9 significantly reduced cell proliferation and 3D colony formation and induced apoptosis. *lnc-HLX-2-7-deleted* D425 Med cells injected into mouse cerebella produced smaller tumors than those derived from parental cells. Pathway analysis revealed that *lnc-HLX2-7* modulated oxidative phosphorylation, mitochondrial dysfunction, and sirtuin signaling pathways. The *MYC* oncogene regulated *lnc-HLX-2-7*, and the small molecule BET-bromodomain (BRD4) inhibitor JQ1 reduced *lnc-HLX2-7* expression.

**Conclusions:** *lnc-HLX-2-7* is oncogenic in MB and represents a promising novel molecular marker and a potential therapeutic target in group 3 MBs in children.

**Key points:**

- *lnc-HLX-2-7* is highly upregulated in group 3 medulloblastomas compared to other sub-groups.
- *In vitro* and *in vivo* studies strongly support an oncogenic role for *lnc-HLX2-7* in group 3 medulloblastoma.
- *lnc-HLX-2-7* may be a novel biomarker and a potential therapeutic target in group 3 medulloblastoma.

**Importance of the study:** Group 3 medulloblastomas are associated with poor clinical outcomes, are difficult to subtype clinically, and their biology is poorly understood. In an effort to address these problems, we identified a group 3-specific long non-coding RNA, *lnc-HLX-2-7*, in an *in silico* analysis of 175 medulloblastomas and confirmed its expression in group 3 medulloblastoma cell lines, patient-derived xenografts, and FFPE samples. CRISPR/Cas9 deletion and antisense oligonucleotide knockdown of *lnc-HLX-2-7* significantly reduced cell growth and 3D colony formation and induced apoptosis. Deletion of *lnc-HLX-2-7* in cells injected into mouse cerebellums reduced tumor growth compared to parental cells, and RNA sequencing of these tumors revealed *lnc-HLX-2-7*-associated modulation of cell viability and cell death signaling pathways. The oncogene *MYC* regulates *lnc-HLX-2-7*, and its expression can be controlled by the BET-bromodomain (BRD4) inhibitor JQ1. *lnc-HLX-2-7* is a candidate biomarker and a potential therapeutic target in group 3 medulloblastomas in children.

## Introduction

Medulloblastoma (MB) is the most common malignant pediatric brain tumor.^1^ Recent large-scale and high-throughput analyses have subclassified MBs into four molecularly distinct subgroups, each characterized by specific developmental origins, molecular features, and prognoses.^1–4^ The well characterized WNT (group 1) and SHH (group 2) subgroups have been causally linked to actived wingless and sonic hedgehog developmental cascades, respectively.^1^ However, significant gaps remain in our understanding of the signaling pathways underlying group 3 and group 4 MBs, which account for 60% of all diagnoses and are frequently metastatic at presentation (~40%).^4^ Group 3 MBs have the worst outcomes and are broadly characterized by a MYC activation signature.^1^ Group 3 and group 4 tumors display significant clinical and genetic overlap, including similar location and presence of isochromosome 17q, and identifying these subgroups can be challenging without the application of multi-gene expression or methylation profiling. Therefore, improved understanding of group 3 tumor drivers and theranostic targets is urgently needed.

Given rapid developments in genome and transcriptome sequencing technologies and the implementation of genomics consortia such as ENCODE and FANTOM, the classical, mRNA-centric view of the transcriptomic landscape has undergone fundamental changes^5^. The vast majority of the genome serves as a template not only for coding RNAs but also non-coding RNAs (ncRNAs). Of the non-coding RNAs, long non-coding RNAs (lncRNAs), which describe a class of RNAs >200 nucleotides in length, have been widely investigated and identified as key regulators of various biological processes including cellular proliferation, differentiation, apoptosis, migration, and invasion.^6–9^ LncRNAs are functionally diverse and participate in transcriptional silencing,^10^ function as enhancers by regulating three-dimensional (3D) chromosomal structure to strengthen interactions between enhancers and promoters,^11^ and sequester miRNAs from their target sites.^12^ LncRNAs can also act as hubs for protein-protein and protein-nucleic acid interactions.^13^ There is now a considerable body of evidence implicating lncRNAs in both health and disease, not least human tumorigenesis.^9,14–19^ It has recently been reported that various lncRNAs play important roles in MB biology,^2,20–23^ although the functional significance of many remains uncertain. Since many lncRNAs are uniquely expressed in specific cancer types,^24^ they may function as powerful MB subgroup-specific biomarkers and therapeutic targets.

By analyzing RNA sequencing data derived from human MBs, here we report that the novel lncRNA *lnc-HLX-2-7* differentiates group 3 from other MBs and normal cerebellum. These *in silico* results were further confirmed by RNA fluorescence *in situ* hybridization (FISH) and qRT-PCR analysis. CRISPR/Cas9 deletion of *lnc-HLX-2-7* in group 3 MB cells significantly reduced cell growth and 3D colony formation and induced apoptosis. Intracranial injection of *lnc-HLX-2-7*-deleted MB cells into mouse cerebellums produced smaller tumors compared to parental cells, and RNA sequencing of xenografts revealed *lnc-HLX-2-7*-associated modulation of cell viability and cell death signaling pathways. *lnc-HLX-2-7* is a promising novel biomarker and potential therapeutic target for group 3 MBs.

## Materials and Methods

### Patient tissue samples

Eighty MB tissue samples obtained from a tumor database maintained by the Department of Pathology at the Johns Hopkins Hospital (JHH) were analyzed (**Supplementary Table 1**) under IRB approved protocol NA_00015113.

### Patient *in silico* data

Raw FASTQ files for RNA sequencing data corresponding to 175 MB patients (referred to as the ICGC dataset) belonging to the four MB subgroups (accession number EGAS00001000215) were downloaded from the European Genome-Phenome Archive (EGA, http://www.ebi.ac.uk/ega/) after obtaining Institutional Review Board approval.^25^

### RNA samples

RNA samples were isolated from normal human cerebellum (BioChain, Newark, CA), MB cell lines and patient-derived xenografts (PDXs). The cell lines DAOY, ONS76, D283 Med, D341 Med, D458 Med, MB002, and HD-MB03 were maintained in the Wechsler-Reya and Raabe labs. The PDXs DMB006, DMB012, RCMB28, RCMB32, RCMB38, RCMB40, RCMB45, and RCMB51 were established in the Wechsler-Reya lab; MED211FH, Med511FH, and MED1712FH were established in the J. Olson lab at Fred Hutchinson Cancer Research Center; BT-084 was created in the T. Milde lab at the German Cancer Research Center (DKFZ) and MB002 was created by Y.J. Cho lab at Oregon Health and Sciences University; all PDXs were maintained in the Wechsler-Reya lab. Functional studies were carried out using D425 Med and MED211 cells maintained in the Eberhart and Raabe labs. CHLA-01 and CHLA-01R were purchased from the American Type Culture Collection (ATCC; Manassas, VA).

### Cell culture

Cell lines were authenticated using single tandem repeat profiling. D425 Med cells were cultured in DMEM/F12 with 10% serum and 1% glutamate/penicillin/streptomycin. MED211 cells were cultured in medium composed of 30% Ham’s F12/70% DMEM, 1% antibiotic antimycotic, 20% B27 supplement, 5 ug/mL heparin, 20 ng/mL EGF, and 20 ng/mL FGF2. DAOY cells were cultured in DMEM with 10% serum and 1% glutamate/penicillin/streptomycin. All cells were grown in a humidified incubator at 37°C, 5% CO_2_.

### Quantitative real-time PCR (qRT-PCR)

Total RNA was purified using the Direct-zol RNA Miniprep kit (Zymo Research, Irvine, CA). To obtain RNA from xenografts, tumor tissues were pulverized and then used for purification. Quantitative PCR was carried out using SYBR Green mRNA assays as previously described.^9^ Primer sequences are listed in **Supplementary Table 2**.

### ASO-*lnc-HLX2-7*

Antisense oligonucleotides (ASOs) were designed using the Integrated DNA Technologies (IDT) Antisense Design Tool (IDT, Coralville, IA). ASO knockdowns were performed with 50 nM (final concentration) locked nucleic acid (LNA) GapmeRs transfected with Lipofectamine 3000 (Thermo Fisher Scientific, Waltham, MA). All ASOs were modified with phosphorothioate (PS) linkages. The following ASOs were used: ASO targeting *lnc-HLX-2-7* (ASO-*lnc-HLX-2-7*): +T*+G*+A*G*A*G*A*T*T*A*A*T*C*T*A*G*A*T*+T*+G*+C and control ASO targeting *luciferase* (*ASO-Luc*): +T*+C*+G*A*A*G*T*A*C*T*C*A*G*C*G*T*A*A*+G*+T*+T. The PS linkages are indicated with * and LNA-modified oligonucleotides are indicated with +.

### siRNA-mediated knockdown of *HLX, MYC*, and *MYCN*

siRNAs targeting *HLX* (catalog no. 4427037, ID: s6639) and *MYC* (catalog no. 4427037, ID: s9129) were purchased from Thermo Fisher Scientific. siRNAs were transfected at 20 nM for 48 h using Lipofectamine RNAiMAX (Thermo Fisher Scientific). The efficiency was determined by qRT-PCR.

### Cell proliferation, apoptosis, and 3D colony formation assays

Cells were plated in 96-well plates at 5 × 10^3^ cells per well in triplicate. After 72 hours of ASO or siRNA transfection, living cells were counted by trypan blue staining. Apoptotic cells were analyzed using a GloMax luminometer (Promega, Fitchburg, WI) with conditions optimized for the Caspase-Glo 3/7 Assay. For the 3D colony formation assay, cells were seeded in 24-well plates at a density of 1 × 10^2^ cells/well and were stained with crystal violet solution approximately 14 days later. Colony number was determined using the EVE cell counter (Nano Entek, Pleasanton, CA), and staining intensity was analyzed using ImageJ software.

### *lnc-HLX-2-7* CRISPR/Cas9 knockdown in D425 Med cells

The single guide RNA (sgRNA) targeting *lnc-HLX-2-7* was designed using Zhang Lab resources (http://crispr.mit.edu/) and synthesized to make the lenti-*lnc-HLX-2-7*-sgRNA-Cas9 constructs as described previously.^26^ The DNA sequences for generating sgRNA were forward: 5’-GGACCCACTCTCCAACGCAG −3’ and reverse: 5’-GCAGGGACCCCTCATTGACG −3’. For the control plasmid, no sgRNA sequence was inserted into the construct. *Lnc-HLX-2-7*-edited cells and control cells were selected using 4 μg/ml puromycin. To determine the genome editing effect, total RNA was extracted from the *lnc-HLX-2-7*-edited cells and control cells and the expression of *lnc-HLX-2-7* quantified by qRT-PCR.

### Medulloblastoma xenografts (intracranial)

All mouse studies were approved and performed in accordance with the policies and regulations of the Animal Care and Use Committee of Johns Hopkins University. Intracranial MB xenografts were established by injecting D425 Med cells and D425 Med cells with *lnc-HLX-2-7* deleted into the cerebellums of NOD-SCID mice (Jackson Laboratory, Bar Harbor, ME). Cerebellar coordinates were −2 mm from lambda, +1 mm laterally, and 1.5 mm deep. Tumor growth was evaluated by weekly bioluminescence imaging using an *in vivo* spectral imaging system (IVIS Lumina II, Xenogen, Alameda, CA).

### Immunohistochemistry

Xenograft tumors were harvested and paraffin-embedded, then tumor sections were stained with hematoxylin and eosin. For the analysis of cell proliferation, tumor sections were incubated with anti-Ki67 (Alexa Fluor 488 Conjugate) antibodies (#11882, 1:200, Cell Signaling Technology, Danvers, MA) at 4°C overnight. For the analysis of apoptosis, DeadEnd™ Fluorometric TUNEL System (Promega) was performed on the tumor sections, according to the manufacturer’s instructions. The stained sections were imaged using a confocal laser-scanning microscope (Nikon C1 confocal system, Nikon Corp, Tokyo, Japan). The acquired images were processed using the NIS (Nikon) and analyzed with the Image J software (https://imagej.nih.gov/ij/). Two tumors (10 fields per tumor section) from each group were analyzed.

### Chromatin immunoprecipitation (ChIP)

ChIP assays were performed based on a modification of previously published methods.^12,27^ Cells (1 × 10^6^) were treated with 1% formaldehyde for 8 minutes to crosslink histones to DNA. The cell pellets were resuspended in lysis buffer (1% SDS, 10 mmol/L EDTA, 50 mmol/L Tris-HCl pH 8.1, and protease inhibitor) and sonicated using a Covaris S220 system (Covaris Inc., Woburn, MA). After diluting the cell lysate 1:10 with dilution buffer (1% Triton-X, 2 mmol/L EDTA, 150 mmol/L NaCl, 20 mmol/L Tris-HCl pH 8.1), diluted cell lysates were incubated for 16 h at 4°C with Dynabeads Protein G (100-03D, Thermo Fisher Scientific) precoated with 5 μL of anti-MYC antibody (ab32, Abcam, Cambridge, UK). ChIP products were analyzed by SYBR Green ChIP-qPCR using the primers listed in **Supplementary Table 2**.

### RNA library construction and sequencing

Total RNA was prepared from cell lines and orthotopic xenografts using Direct-zol RNA Miniprep kits (Zymo Research, Irvine, CA). RNA quality was determined with the Agilent 2100 Bioanalyzer Nano Assay (Agilent Technologies, Santa Clara, CA). Using a TruSeq Stranded Total RNA library preparation Gold kit (Illumina Inc., San Diego, CA), strand-specific RNA-seq libraries were constructed as per the instructions. The quantification and quality of final libraries were determined using KAPA PCR (Kapa Biosystems, Waltham, MA) and a high-sensitivity DNA chip (Agilent Technologies), respectively. Libraries were sequenced on an Illumina NovaSeq 6000 using 1 × 50 base paired-end reads.

### Single-cell RNA-seq library construction and sequencing

Cell suspensions required for generating 8000 single cell gel beads in emulsion (GEM) were loaded onto the Chromium controller (10X Genomics, Pleasanton, CA). Each sample was loaded onto the single cell 3’ v3.1 chip. The 3’ gene expression library was prepared using a Chromium v3.1 single cell 3’ library kit (10X Genomics). The quantification and quality of final libraries were determined using a KAPA PCR (Kapa Biosystems) and a high sensitivity DNA chip (Agilent Technologies), respectively. Samples were diluted to 1.8 pM for loading onto the NextSeq 550 (Illumina) with a 150-cycle paired-end kit using the following read length: 28 cycles for Read 1, 8 cycles i7 index, 0 cycles i5 index, and 91 cycles Read 2. Detailed methods of sequence and data analysis are described in **Supplementary Methods.**

### Ingenuity pathway analysis (IPA)

To analyze pathways affected by *lnc-HLX-2-7*, differentially expressed genes between D425 Med and D425 Med with *lnc-HLX-2-7* deleted were compiled and analyzed using Qiagen’s IPA. Analysis was conducted via the IPA web portal (www.ingenuity.com).

### Data availability

The organoid data described in the manuscript is accessible at NCBI GEO accession number GSE151810.

### RNA-fluorescence *in situ* hybridization (RNA-FISH)

RNA was visualized in paraffin-embedded tissue sections using the QuantiGene ViewRNA ISH Tissue Assay Kit (Affymetrix, Frederick, MD). In brief, tissue sections were rehydrated and incubated with proteinase K. Subsequently, they were incubated with ViewRNA probesets designed against human *lnc-HLX-2-7, MYC*, and *MYCN* (Affymetrix, Santa Clara, CA). Custom Type 1 primary probes targeting *lnc-HLX-2-7*, Type 6 primary probes targeting *MYC*, and Type 6 primary probes targeting *MYCN* were designed and synthesized by Affymetrix (**Supplementary Table 2**). Hybridization was performed according to the manufacturer’s instructions.

### Statistical analysis

Statistical analyses were performed using GraphPad Prism software and Limma R package. Data are presented as mean ± SD of three independent experiments. Differences between two groups were analyzed by the paired Student’s *t*-test. Kruskal–Wallis analysis was used to evaluate the differences between more than two groups. Survival analysis was performed using the Kaplan–Meier method and compared using the log-rank test.

## Results

### Identification of the group 3-specific long-noncoding RNA, *lnc-HLX-2-7*

To identify MB group 3-specific lncRNAs, we obtained 175 RNA-seq files (FASTq) representing the four MB subgroups (WNT, SHH, group 3 and group 4) from the European Genome-Phenome Archive (EGA) and applied combined GENCODE and LNCipedia annotations.^28^ Given the need to find novel biomarkers that differentiate group 3 from other groups, we identified a set of lncRNAs (*lnc-HLX-1, lnc-HLX-2, lnc-HLX-5*, and *lnc-HLX-6*) with markedly elevated expression in group 3 MB (**Fig. 1A and Supplementary Table 3**). *lnc-HLX-1, lnc-HLX-2, lnc-HLX-5* and *lnc-HLX-6* showed a high expression correlation (**Fig. 1B**) and were highly expressed in group 3 MB patient samples compared to other subgroups (*p*<0.01, **Fig. 1C**). We recently reported that some of these lncRNAs also show group 3-specific differential expression.^29^ Due to *lnc-HLX-2*’s proximity to its host coding gene transcription factor and homeobox gene *HB24* (*HLX*) and a recent study reporting that the *lnc-HLX-2* region is a group 3 MB-specific enhancer region (**Supplementary Fig. 1**),^30^ we focused on *lnc-HLX-2. lnc-HLX-2* is located 2300 bp downstream of the transcriptional start site (TSS) of *HLX* (**Supplementary Fig. 2A**) and consists of 11 transcripts (*lnc-HLX-2-1* to *lnc-HLX-2-11*; **Supplementary Fig. 2B**), of which *lnc-HLX-2-7* was highly expressed in group 3 MBs (**Supplementary Fig. 2C**). qRT-PCR analysis verified that *lnc-HLX-2-7* was highly upregulated in group 3 MB cell lines (**Fig. 1D**) and xenografts (**Fig. 1E**) compared to other groups.

**Figure 1.**
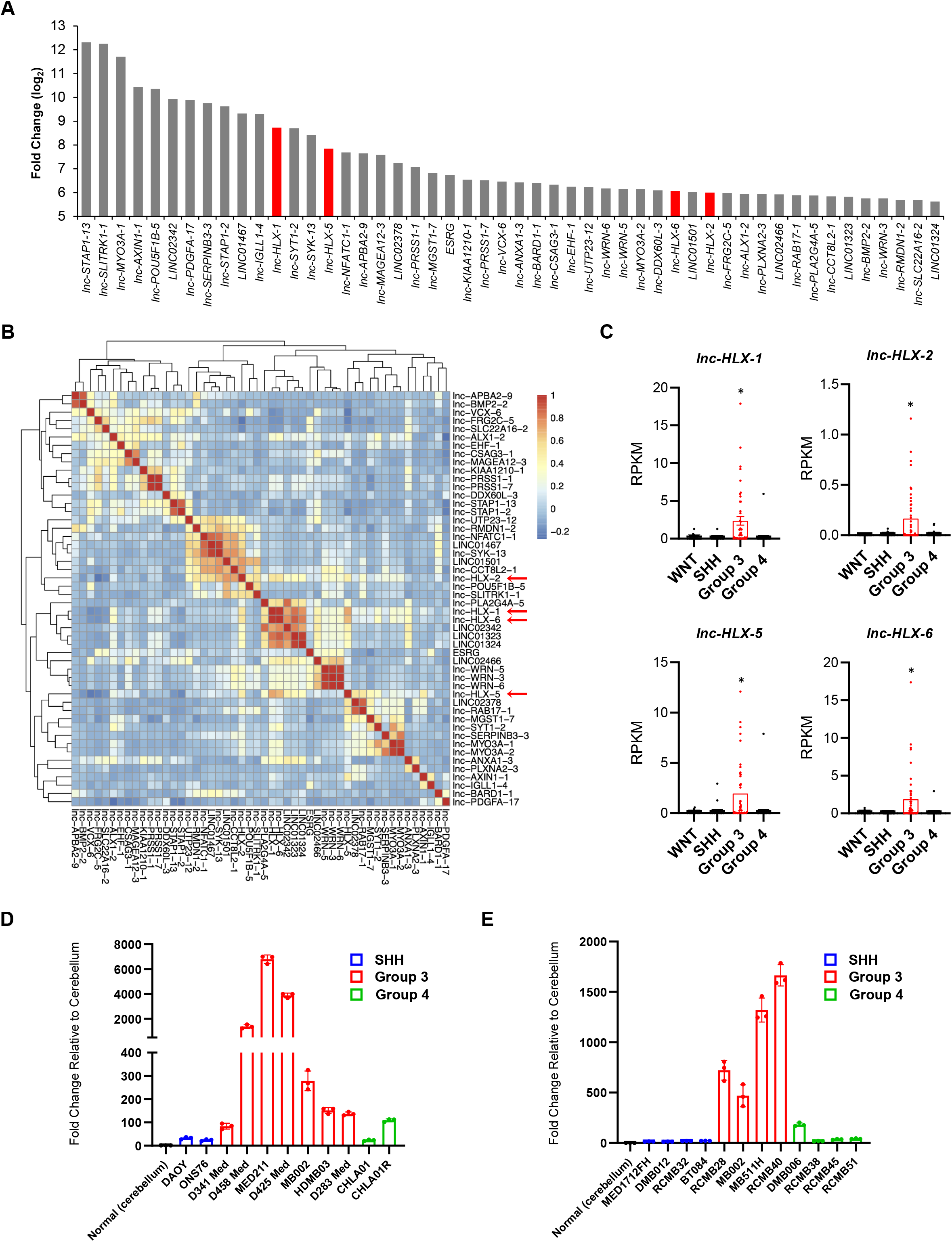
Identification and validation of group 3-specific lncRNA, *lnc-HLX-2-7*. (A) Top 50 lncRNAs with the highest expression in group 3 MBs compared to other MB sub-groups are shown. y-axis indicates log2-fold change value of each lncRNA. (B) The heat map represents the similarity of expression within group 3 MBs of each lncRNA shown in (A). (C) Boxplot showing distribution of normalized expression values of *lnc-HLX-1*, *lnc-HLX-2*, *lnc-HLX-5*, and *lnc-HLX-6* in WNT, SHH, group 3 and group 4 MBs. Dots represent the expression value for each MB patient. \**p* < 0.01, Kruskal–Wallis analysis. (D, E) qRT-PCR analysis showing the distribution of normalized expression values of *lnc-HLX-2-7* in MB cell lines (D) and PDX samples (E) of group 3, group 4, and SSH MBs. Values indicate fold change relative to cerebellum.

### *lnc-HLX-2-7* functions as an oncogene *in vitro*

To investigate the function of *lnc-HLX-2-7*, we used antisense oligonucleotides (ASOs) to inhibit *lnc-HLX-2-7* expression in MED211 and D425 Med MB cells. Transfection with *ASO-lnc-HLX-2-7* significantly decreased *lnc-HLX-2-7* expression compared to controls (ASO-luc) in both cell lines (*p*<0.01, **Fig. 2A**), which significantly suppressed MB cell growth and induced apoptosis (*p*<0.01, **Fig. 2B and 2C**). Next, CRISPR/Cas9 knock-down was used to generate single-cell colonies and further investigate the effect of *lnc-HLX-2-7* in MB cells. We generated stable D425 Med-*lnc-HLX-2-7*-sgRNA cells, which constitutively expressed sgRNAs against *lnc-HLX-2-7* to lower the *lnc-HLX-2-7* expression (**Fig. 2D**). As expected, D425 Med-*lnc-HLX-2-7*-sgRNA cells showed a decrease in growth (**Fig. 2E**) and reduced colony-forming ability (**Fig. 2F**) when compared D425 Med control cells *in vitro*.

**Figure 2.**
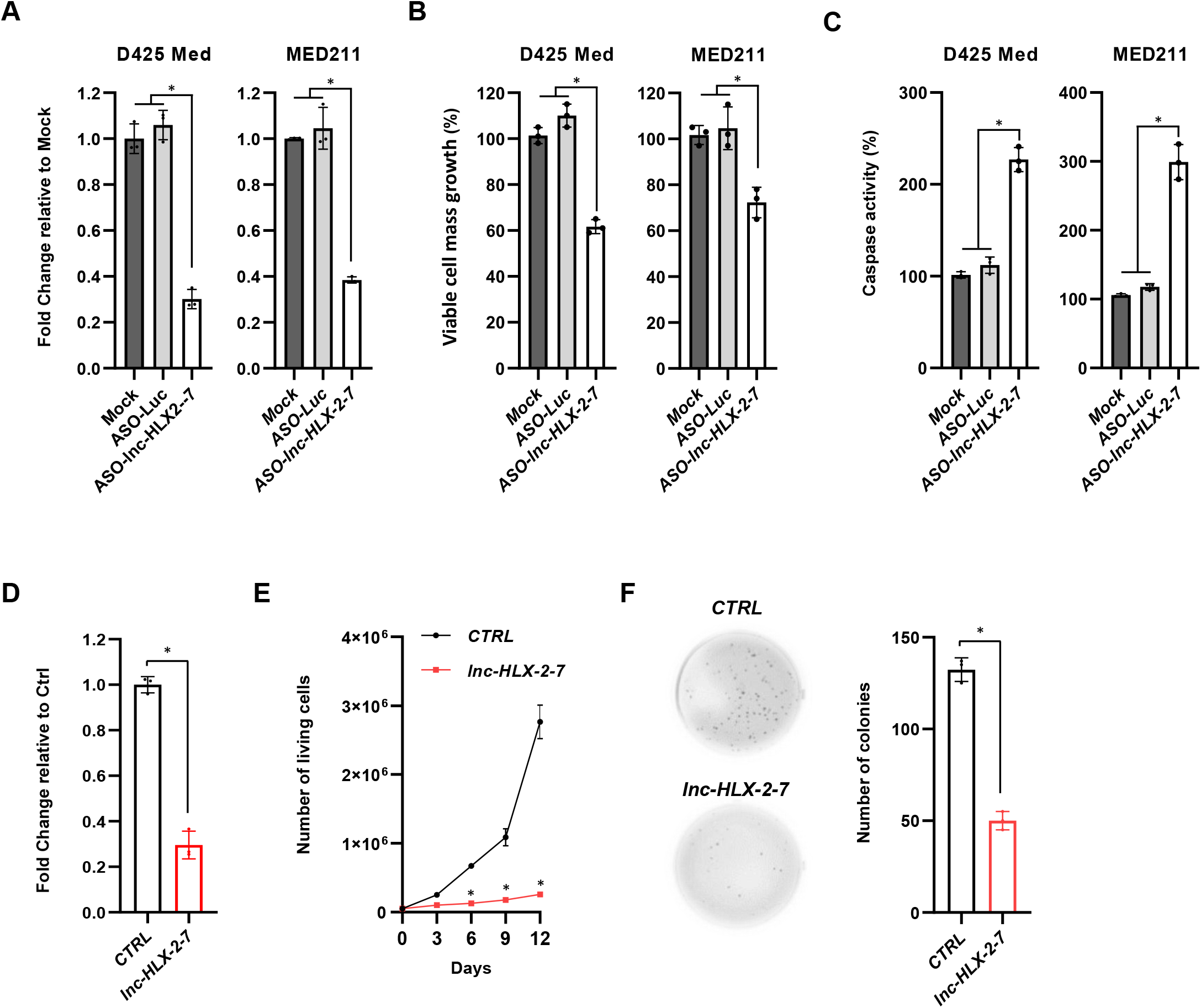
Effects of *lnc-HLX-2-7* expression on the proliferation and apoptosis of group 3 MB cells. (A) Expression level of *lnc-HLX-2-7* in D425 Med and MED211 cells treated with ASO against the genes indicated on the x-axis. Relative expression level to mock (non-transfected) is indicated on the y-axis. \**p* < 0.01, Kruskal–Wallis analysis. Viable cell numbers (B) and apoptotic cell numbers (C) in D425 Med and MED211 cells treated with either ASO-luc or ASO-*lnc-HLX-2-7*. Relative value to mock is indicated on the y-axis. \**p* < 0.01, Kruskal-Wallis analysis. (D) Expression level of *lnc-HLX-2-7* in D425 Med control (*CTRL*) and D425-*lnc-HLX-2-7*-sgRNA (lnc-HLX-2-7) cells. Relative expression level to *CTRL* is indicated on the y-axis. \**p* < 0.01, Student’s *t*-test. (E) Cell viability assays performed with D425 Med control (*CTRL*) and D425 Med-*lnc-HLX-2-7*-sgRNA (*lnc-HLX-2-7*) cells. Points represent the mean and standard deviation of three biological replicates. \**p* < 0.01, Student’s *t*-test. (F) Colony formation assays performed with D425 Med control (*CTRL*) and D425 Med-*lnc-HLX-2-7*-sgRNA (*lnc-HLX-2-7*) cells. Three independent experiments were performed, and data are presented as mean ± SD. \**p* < 0.01, Student’s *t*-test.

While the functions of the majority of lncRNAs are not yet known, some have been shown to function *in cis* by regulating the expression of neighboring genes.^31–33^ Since *lnc-HLX-2-7* is located downstream of the *HLX* transcription start site (TSS; **Supplementary Fig. 2A**), we determined whether *lnc-HLX-2-7* regulates *HLX* expression; indeed, *HLX* expression was significantly reduced in D425 and MED211 cells following treatment with *ASO-lnc-HLX-2-7* (**Supplementary Fig. 3**). In addition, *HLX* knockdown significantly decreased the growth of D425 Med and MED211 cells (**Supplementary Fig. 4**). While the current study focuses on the role of lncRNA *HLX-2-7*, understanding the molecular function of its host-coding gene *HLX* requires further investigation, which is ongoing.

### *lnc-HLX-2-7* regulates tumor formation in mouse intracranial xenografts

To evaluate the effect of *lnc-HLX-2-7* on tumor growth *in vivo*, we established intracranial MB xenografts in NOD-SCID mice. D425 Med control cells and D425 Med-*lnc-HLX-2-7*-sgRNA cells were pre-infected with a lentivirus containing a luciferase reporter. Weekly evaluation of tumor growth by bioluminescence imaging revealed significantly smaller tumors in mice transplanted with D425 Med-*lnc-HLX-2-7*-sgRNA cells compared to mice transplanted with control cells (n=3, *p*<0.05, **Fig. 3A and 3B**). At day 35, tumors were harvested and cut into sections and then subjected to Ki67 and TUNEL staining. Ki67 analysis showed reduced cell proliferation in D425 Med-*lnc-HLX-2-7*-sgRNA cell-transplanted mice (*p*<0.01, **Fig. 3C**). TUNEL analysis found out that *lnc-HLX-2-7* depletion induced significantly higher percentage of TUNEL-positive cells than compared to mice transplanted with control cells (*p*<0.01, **Fig. 3D**). Kaplan-Meier plots demonstrated that the group transplanted with D425 Med-*lnc-HLX-2-7*-sgRNA cells had significantly prolonged survival compared to the control (**Fig. 3E**). Together, these results demonstrate that *lnc-HLX-2-7* regulates tumor growth *in vivo* and may function as an oncogene.

**Figure 3.**
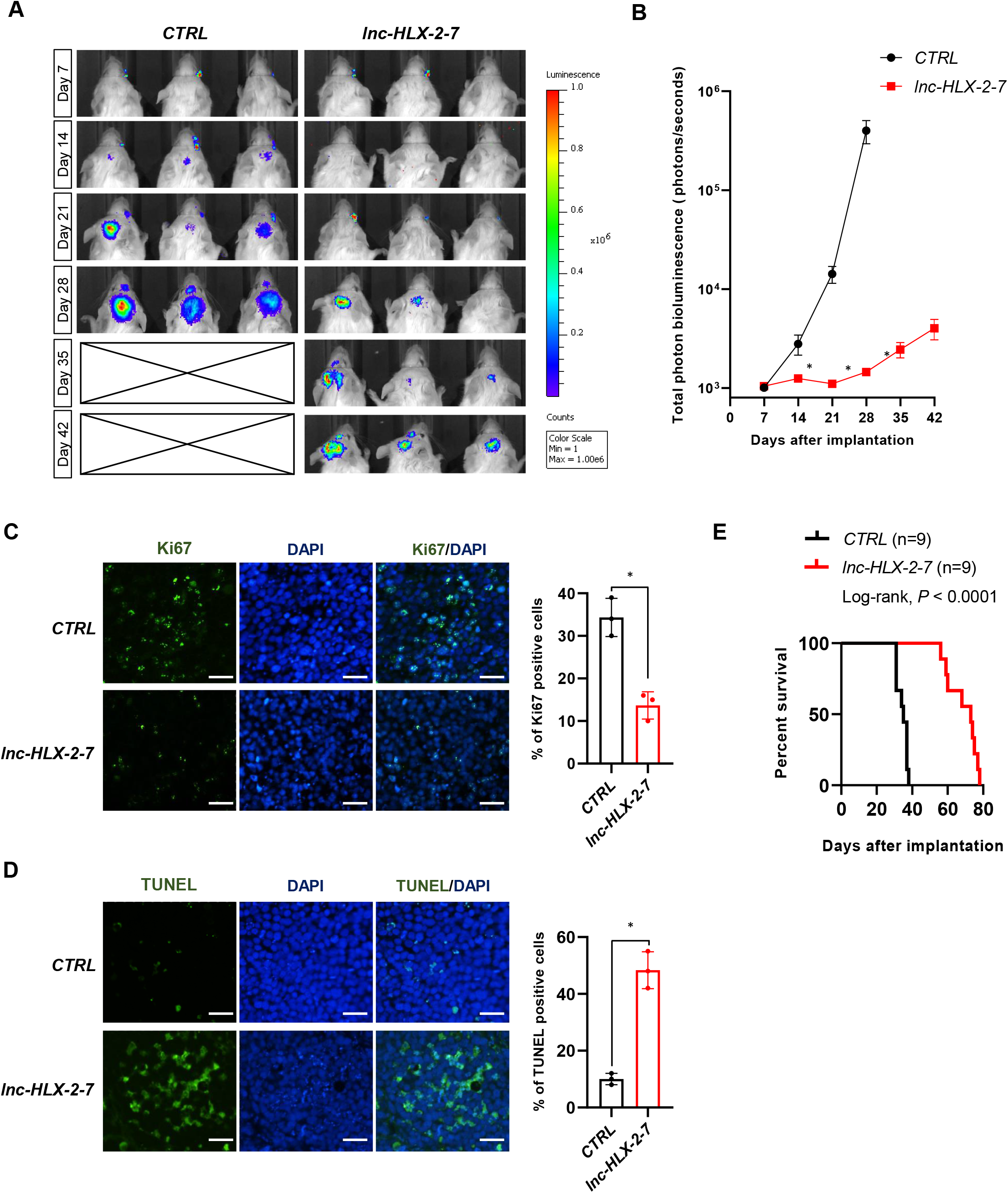
*lnc-HLX-2-7* promotes the tumorigenicity of group 3 MB cells *in vivo*. (A) D425 Med control (CTRL) and D425 Med-lnc-HLX-2-7-sgRNA (lnc-HLX-2-7) cells, expressing luciferasewere implanted into the right forebrain of NOD-SCID mice, and tumor formation was assessed by bioluminescence imaging. Changes in bioluminescent signal were examined weekly after tumor implantation. (B) Quantification of total photon counts from mice implanted with D425 Med control (*CTRL*) and D425 Med-lnc-HLX-2-7-sgRNA (*lnc-HLX-2-7*) cells. n=3, \**p* < 0.05, Student’s *t*-test. (C) Ki67 and (D) TUNEL staining of xenograft tumors. Nuclei are stained with DAPI. Scale bars, 50 μm. Quantification of Ki67 and TUNEL-positive cells were shown. \**p* < 0.05, Student’s *t*-test. (E) Overall survival was determined by Kaplan-Meier analysis, and the log-rank test was applied to assess the differences between groups.

### Transcriptional regulation of *lnc-HLX-2-7* by the *MYC* oncogene

Since the majority of group 3 tumors exhibit elevated expression and amplification of the *MYC* oncogene,^2,34^ we hypothesized that MYC may regulate the expression of *lnc-HLX-2-7*. We therefore knocked down *MYC* by siRNA in D425 Med cells, which decreased the expression of both*MYC* and *lnc-HLX-2-7* (**Fig. 4A**), suggesting that MYC may be an upstream regulator (direct or indirect) of *lnc-HLX-2-7*. To further support this, we also identified a MYC-binding motif (E-box; -*CACGTG*-) 772 bp upstream of the putative TSS of *lnc-HLX-2-7* using the JASPAR CORE database (http://jaspar.binf.ku.dk/)^35^ (**Fig. 4B**). To test whether MYC could interact with the endogenous *lnc-HLX-2-7* promoter, chromatin immunoprecipitation (ChIP) was performed in group 3 MB D425 Med cells. DAOY, a SHH MB cell line, was used as negative control, since *lnc-HLX-2-7* is not expressed in SHH MBs. MYC bound to the E-box motif within the upstream region of *lnc-HLX-2-7* in D425 cells, but not in DAOY cells (**Fig. 4C**). These results strongly suggest that MYC is a direct regulator of *lnc-HLX-2-7*.

**Figure 4.**
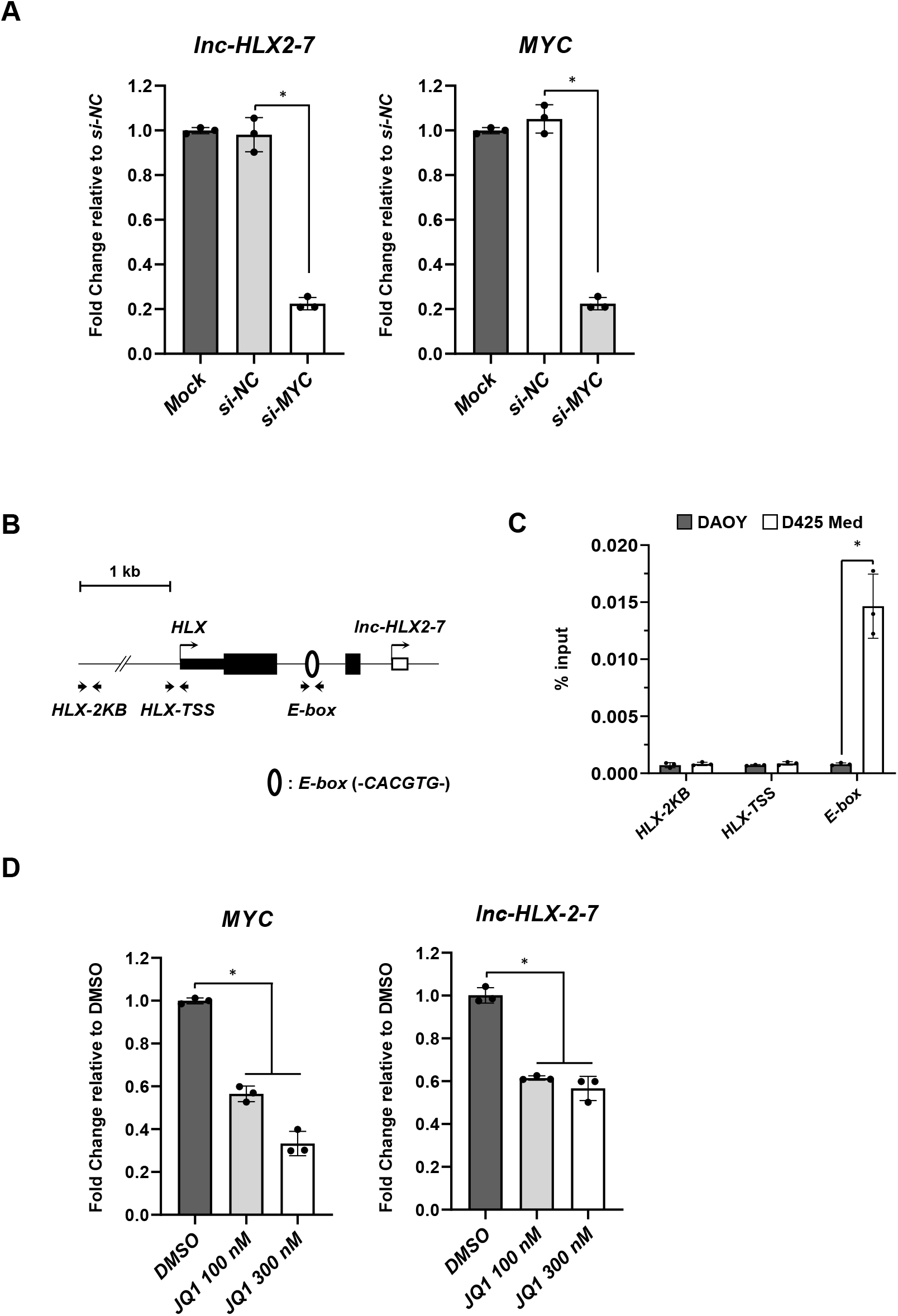
MYC regulates the expression of *lnc-HLX-2-7* in group 3 medulloblastomas. (A) Expression level of *MYC* and *lnc-HLX-2-7* in D425 cells treated with siRNA against the indicated genes on the x-axis. Relative expression level to mock (non-transfected) is indicated on the y-axis. \**p* < 0.01, Kruskal–Wallis analysis. (B) Schematic diagram showing E-box motifs around the TSS of *lnc-HLX-2-7*. Open circles indicate E-box motifs. (C) Enrichment of MYC in the *lnc-HLX-2-7* promoter regions in DAOY and D425 Med cells. Enrichment is expressed as a percentage of input DNA. \**p* < 0.01, Student’s *t*-test. (C) Expression level of *MYC* and *lnc-HLX-2-7* in D425 cells treated with JQ1. Values are indicated relative to abundance in DMSO-treated cells. \**p* < 0.01, Kruskal–Wallis analysis.

### JQ1 regulates *lnc-HLX2-7* via MYC

Several previous studies have demonstrated that BRD4, a member of the bromodomain and extraterminal domain (BET) family, regulates *MYC* transcription and that JQ1 effectively suppresses cancer cell proliferation by inhibiting BRD4-mediated regulation of MYC in various types of cancer including MB^36–40^. To test the JQ1 effect on *lnc-HLX-2-7* regulation, we treated D425 Med cells with different doses (100 or 300 nM) of the drug. As shown in **Fig. 4D**, both *MYC* and *lnc-HLX-2-7* were downregulated in D425 Med cells. Collectively, our results show that BRD4 inhibitors can be used to target MYC-mediated regulation of *lnc-HLX-2-7* expression.

### RNA sequencing detects *lnc-HLX-2-7* interacting genes and pathways in medulloblastoma

To gain further insights into the functional significance of *lnc-HLX-2-7*, gene expression was measured by RNA-seq in D425 Med-*lnc-HLX-2-7*-sgRNA cells and in xenografts derived from them. Among 1033 genes with a significant change in expression (FDR <0.05), 484 genes were upregulated and 549 genes were downregulated in cultured D425 Med-*lnc-HLX-2-7*-sgRNA cells (**Supplementary Fig. 5A**). Ingenuity Pathway Analysis (IPA) revealed that *lnc-HLX-2-7* knockdown preferentially affected genes associated with cell death (**Supplementary Fig. 5B**). Of note, upstream regulator analysis showed that these genes contribute to important cancer pathways including MYC, KRAS, HIF1A, and EGFR signaling (**Supplementary Fig. 5C**).

In D425 Med-*lnc-HLX-2-7*-sgRNA cell-transplanted xenografts, among 540 genes with a significant change in expression (FDR<0.05), 409 genes were upregulated and 131 genes were downregulated (**Fig. 5A**). IPA analysis revealed that *lnc-HLX-2-7* knockdown preferentially regulated genes associated with cell viability, including in neuronal cells (**Fig. 5B**). Canonical IPA pathway analysis showed that oxidative phosphorylation, mitochondrial dysfunction, and sirtuin signaling pathways were highly modulated by *lnc-HLX2-7* (**Fig. 5C**). Ten differentially expressed genes detected by RNA-seq and pathway analysis were validated by qRT-PCR, and the cancer and brain tumor-related genes *PTGR1, FZD6, TRPM3, NAMPT, NRBP2, NBAT1, CCNG2, ELK4, CDKN2C*, and *CDK6* were all dysregulated in *lnc-HLX-2-7-depleted* xenografts (**Supplementary Fig. 6**).

**Figure 5.**
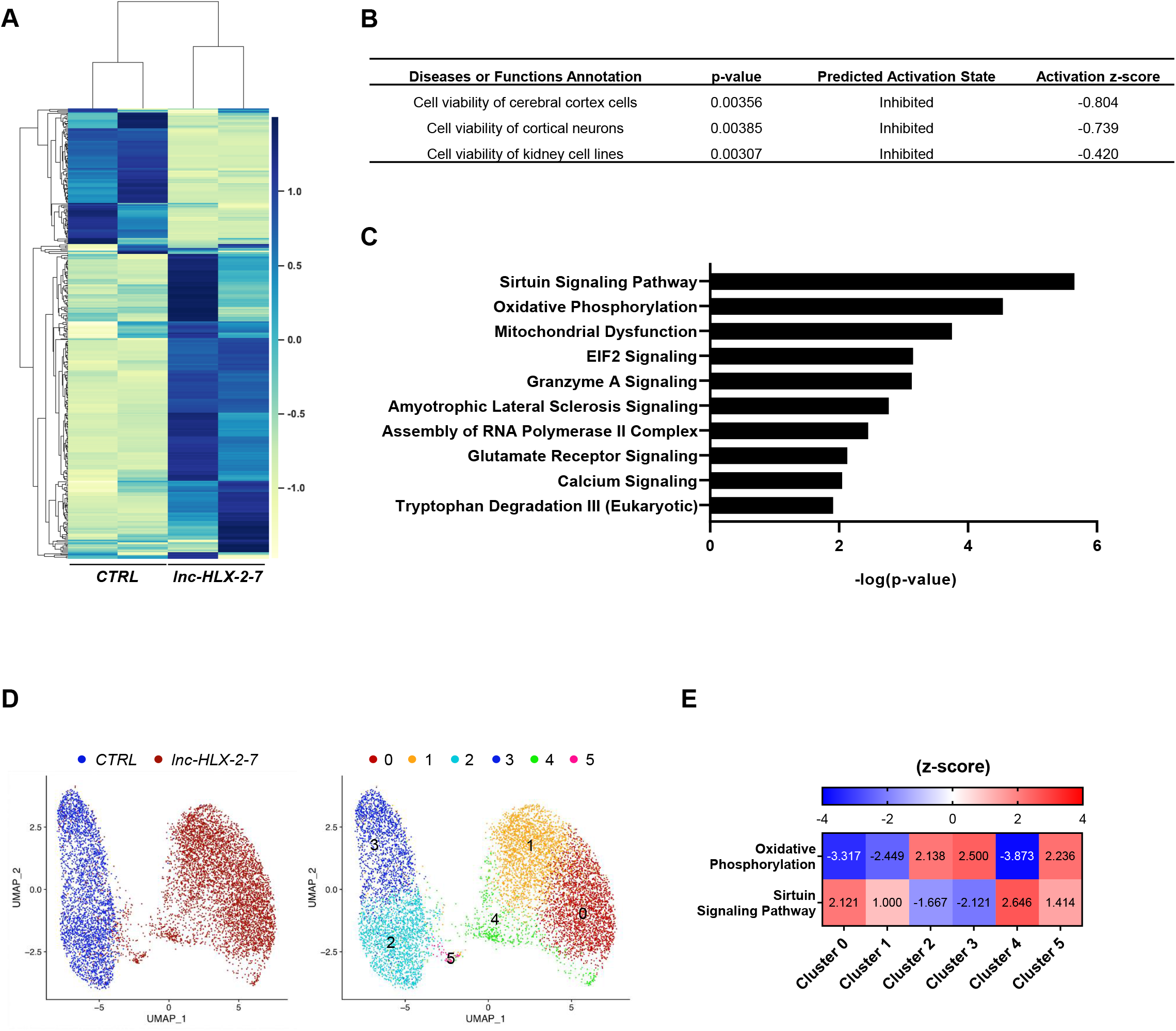
RNA sequencing detects *lnc-HLX-2-7* interacting genes and pathways. (A) Heatmap representation of genes up and downregulated after *lnc-HLX2-7* depletion in D425 xenografts. (B) Molecular and cellular functions and diseases associated with these genes. (C) Canonical IPA analysis was performed to predict signaling pathway activity. The 10 most significant pathways with (with lowest *p*-values) are presented. (D) Uniform Manifold Approximation and Projection (UMAP) visualization of transcriptionally distinct cell populations from D425 Med and *lnc-HLX-2-7-deleted* xenograft samples (3,442 cells from D425 Med and 6,193 cells from *lnc-HLX-2-7-deleted* xenograft). (E) Canonical IPA analysis to predict signaling pathway activity in cluster 0, 1, 2, 3, 4, and 5. The top canonical pathways with (with lowest *p*-values) are presented.

Xenograft tumors were further characterized by single-cell sequencing. Subsequent to quality control, 3,442 and 6,193 cells were obtained for D425 and *lnc-HLX-2-7* deleted D425 respectively. Integrated analysis and clustering of D425 control and *lnc-HLX-2-7* depleted xenografts resulted in 6 clusters of single cells. Clusters 2 and 3 are almost entirely D425 control xenograft. Clusters 0, 1, 4, and 5 are almost exclusively *lnc-HLX-2-7* depleted xenograft (**Fig. 5D**). The top canonical pathways including oxidative phosphorylation and sirtuin signaling pathways were impacted in *lnc-HLX-2-7* depleted single cell populations compared to D425 control (**Fig. 5E**).

### *lnc-HLX-2-7* expression is specific to group 3 MBs

We next confirmed group 3 specificity by visualizing *lnc-HLX-2-7* expression by RNA-FISH in formalin-fixed, paraffin-embedded tissue samples from D425 Med mouse xenografts and patients with MB. *lnc-HLX-2-7* was expressed in D425 Med mouse xenografts but not normal brain (**Supplementary Fig. 7**), and *lnc-HLX-2-7* was readily detected in all group 3 MB samples but not in group 4 MBs (**Fig. 6A and 6B**). Quantitative analysis of the tissues further confirmed significantly higher *lnc-HLX-2-7* expression in group 3 MBs compared to group 4 and SHH MBs (n=20, *p* < 0.01, **Fig. 6C and Supplementary Fig. 8**). Finally, *lnc-HLX-2-7* overexpression was associated with poor patient outcomes in group 3 MB (**Fig. 6D**). Collectively, our analyses suggest that *lnc-HLX-2-7* expression is specific to group 3 MBs and can be detected using an assay readily applicable to the clinical setting.

**Figure 6.**
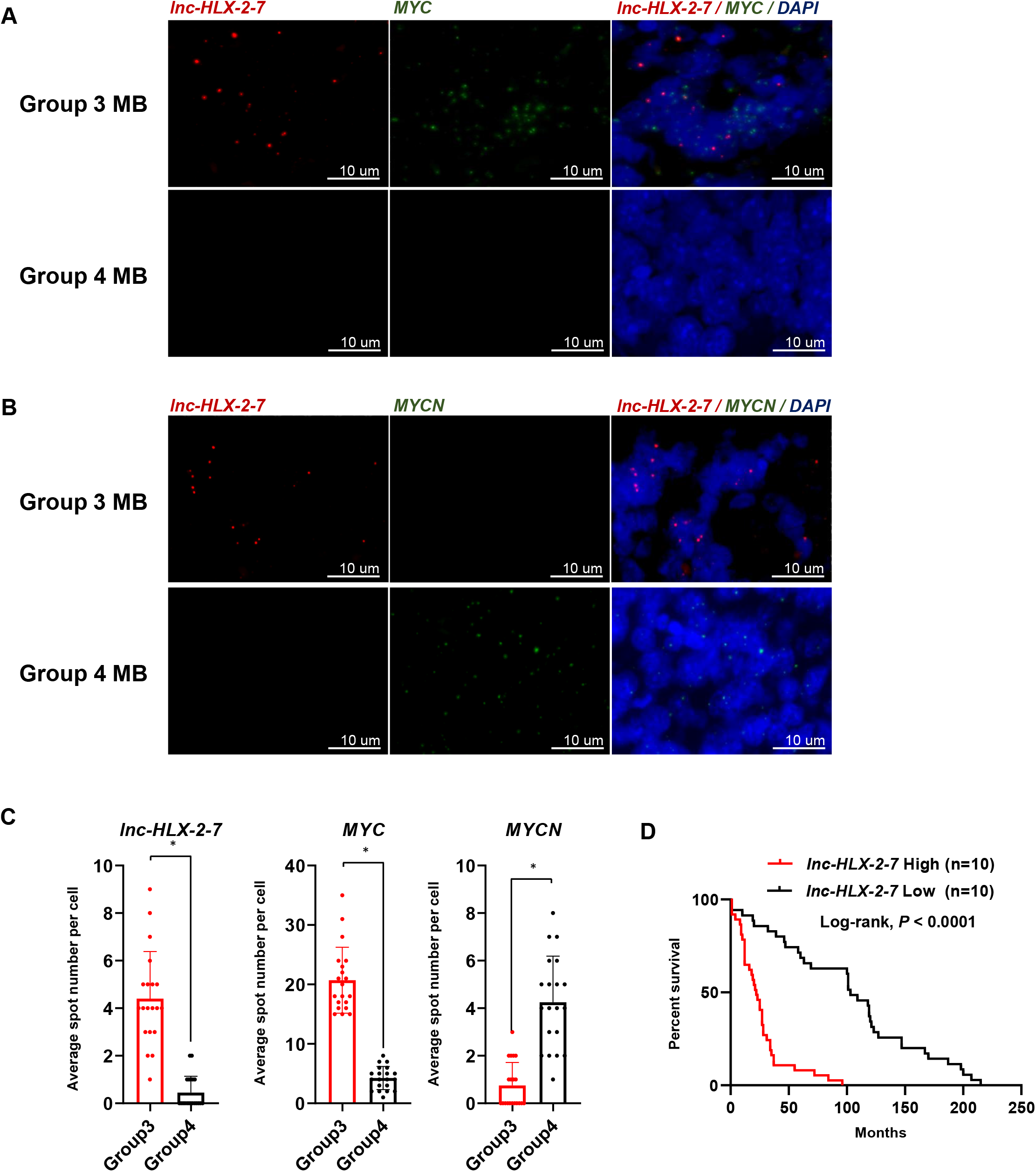
RNA-FISH confirms that *lnc-HLX-2-7* expression is specific to group 3 MB patients. (A) Representative RNA-FISH analysis of *lnc-HLX-2-7* and *MYC* in MB tissues. RNA-FISH analysis of *lnc-HLX-2-7* and *MYC* in group 3 MB patients (upper panels) and group 4 MB patients (lower panels). (B) Representative RNA-FISH analysis of *lnc-HLX-2-7* and *MYCN* in MB tissues. RNA-FISH analysis of *lnc-HLX-2-7* and *MYCN* in group 3 MB patients (upper panels) and group 4 MB patients (lower panels). Nuclei are stained with DAPI. Scale bars, 10 μm. (C) The spot numbers relating to *lnc-HLX-2-7*, *MYC*, and *MYCN* were quantified per cell in group 3 and group 4 MB patients. n=20, \**p* < 0.01, Student’s *t*-test. (D) Kaplan–Meier survival curves of group 3 MB patients according to *lnc-HLX-2-7* expression.

## Discussion

The functions and clinical relevance of lncRNAs in MB are poorly described. Here we provide evidence that the lncRNA *lnc-HLX-2-7* is clinically relevant and biologically functional in group 3 MBs. Using publicly available patient-derived RNA-seq datasets, we discovered that *lnc-HLX-2-7* expression is particularly high in group 3 MBs compared to other groups. By depleting the expression of *lnc-HLX-2-7* by CRISPR/Cas9 and ASOs, we showed both *in vitro* and *in vivo* that *lnc-HLX-2-7* knockdown reduced proliferation and colony formation and increased apoptosis in MB.

The region encoded by *lnc-HLX-2-7* has been reported as an MB-specific enhancer region.^30^ Therefore, ncRNAs transcribed from this region may function as enhancer RNAs (eRNAs), a class of lncRNAs synthesized at enhancers, and may regulate the expression of their surrounding genes. We found that *lnc-HLX-2-7* positively regulated the expression of the adjacent *HLX* gene. Although the mechanism by which *lnc-HLX-2-7* regulates *HLX* remains unclear, *lnc-HLX-2-7* may function as an eRNA in this context. *HLX* has recently been shown to be a key gene mediating BET inhibitor responses and resistance in group 3 MBs.^41^ In this study, we discovered that *lnc-HLX-2-7* controls *HLX* expression and contributes to MB cell proliferation so it is possible that it may influence BET inhibitor resistance. In addition, our results show that the *MYC* oncogene regulates *lnc-HLX-2-7* expression. A recent report suggests that the small molecule JQ1, a BET inhibitor that disrupts interactions with MYC, could be a therapeutic option to treat group 3 MBs.^42^ However, group 3 MB tumors may also become resistant to BET inhibitor through mutations in the *BRD4* gene, and transcription factors like MYC and HLX are poor therapeutic targets with short half-lives and pleiotropic properties.^43^ We postulate that *lnc-HLX-2-7* inhibition may provide a novel solution to BET inhibitor resistance or amplify the effects of BET inhibitors, a hypothesis that requires further investigation.

Recent evidence shows that HLX directly regulates several metabolic genes and controls mitochondrial biogenesis^44^. In the present study, we demonstrate that *lnc-HLX2-7* modulated oxidative phosphorylation, mitochondrial dysfunction, and sirtuin signaling pathways in intracranial xenograft models. These findings suggest that *lnc-HLX-2-7* contributes to the metabolic state of group 3 MBs by regulating HLX expression. This newly discovered link between *lnc-HLX-2-7* and metabolism may have important therapeutic implications.

Group 3 and group 4 MBs display clinical and genetic overlap, with similar anatomic location and presence of isochromosome 17q, so it is not currently possible to distinguish them without applying multi-gene expression or methylation profiling. *lnc-HLX-2-7* may represent a useful single molecular marker that could distinguish group 3 from group 4 MBs. Furthermore, RNA-FISH using probes targeting *lnc-HLX-2-7*, a technique readily applicable in clinical laboratories, readily discriminated group 3 from group 4 MBs.

In conclusion, we show that the lncRNA *lnc-HLX-2-7* is clinically and functionally relevant in group 3 MBs. Future studies will determine the mechanism by which *lnc-HLX-2-7* promotes MB tumorigenesis. Together, our findings support the hypothesis that lncRNAs, and *lnc-HLX-2-7* in particular, are functional in human MBs and may offer promising future opportunities for diagnosis and therapy.

## Acknowledgements

This work was supported by National Institutes of Health grants NCI 5P30CA030199 (SBP), P30 CA006973 (JHU SKCCC), Hough Foundation, and the Schamroth Project funded by Ian’s Friends Foundation to R.J.P. We thank Ms. Tiffany Casey for helping us with the manuscript formatting.

## Notes

**Funding:** Schamroth Project funded by Ian’s Friends Foundation to R.J.P, G.J., and C.B. and the Hough Foundation grant to R.J.P and G.J. This study was also supported by P30 CA006973 (JHU SKCCC) to R.J.P. C.G.E, E.R., and C.B. and NCI 5P30CA030199 (SBP) to R.W-R. and R.J.P.

**Conflicts of interest:** The authors declare no conflicts of interest.

### Competing Interest Statement

The authors have declared no competing interest.

### Summary of Updates

Author affiliations updated Supplemental files updated

