## Supplementary figures for "The long non-coding RNA *lnc-HLX-2-7* is oncogenic in group 3 medulloblastomas"

**Supplementary Figure 1** **Katsushima et al.**

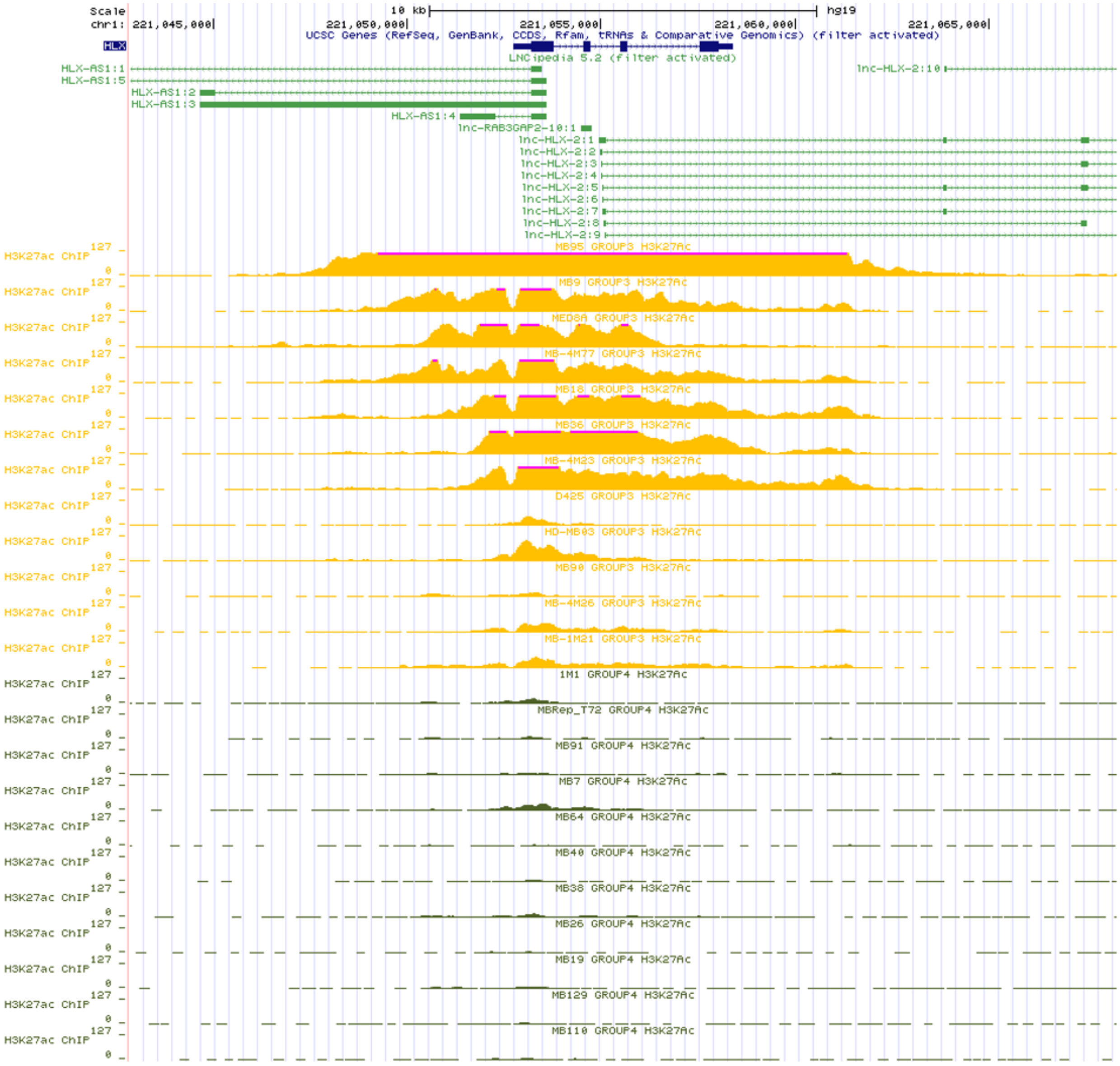


**Supplementary Figure 1. UCSC Genome Browser views showing H3K27ac ChIP-seq plots for *lnc-HLX-2* in group 3 and group 4 MBs.**

**Supplementary Figure 2** **Katsushima et al.**

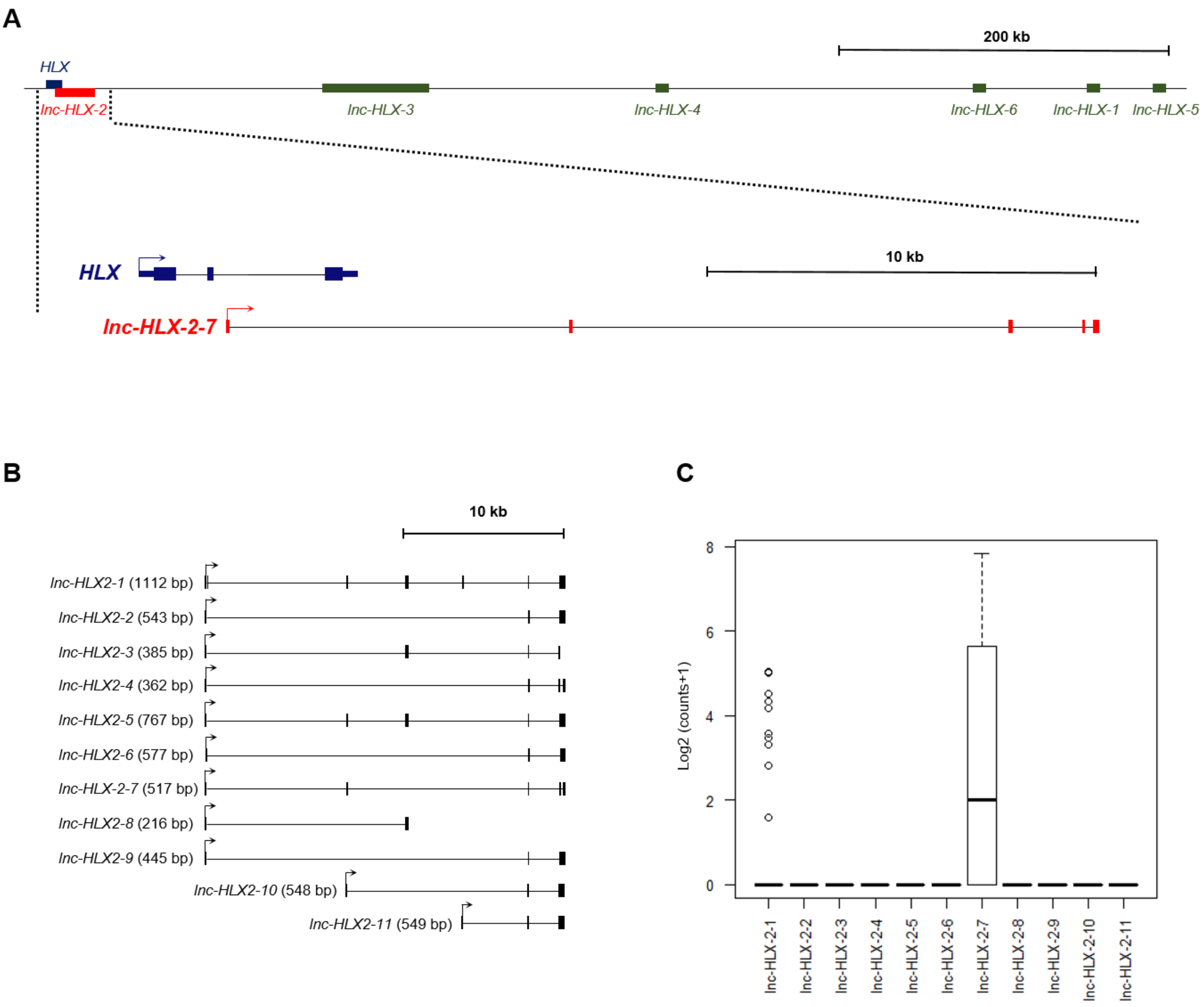

**Supplementary Figure 2. Location of *HLX* and *lnc-HLX-2-7* and expression levels of *lnc-HLX-2-7* variants**

(A) *lnc-HLX-2-7* is a 517bp intronic lncRNA encoded within the *HLX* gene and is located 2300 bp downstream of the *HLX* gene. The fourth and the fifth exon of the *lnc-HLX-2-7* are repeated elements. The first exon has a 32 bp repeat at its end, while second and the third exon are non-repeated. (B) *lnc-HLX-2* contains 11 transcripts (*lnc-HLX-2-1* to *lnc-HLX-2-11*) (C) Boxplot showing distribution of normalized expression values of 11 transcripts (*lnc-HLX-2-1* to *lnc-HLX-2-11*) of *lnc-HLX2* in group 3 MBs.

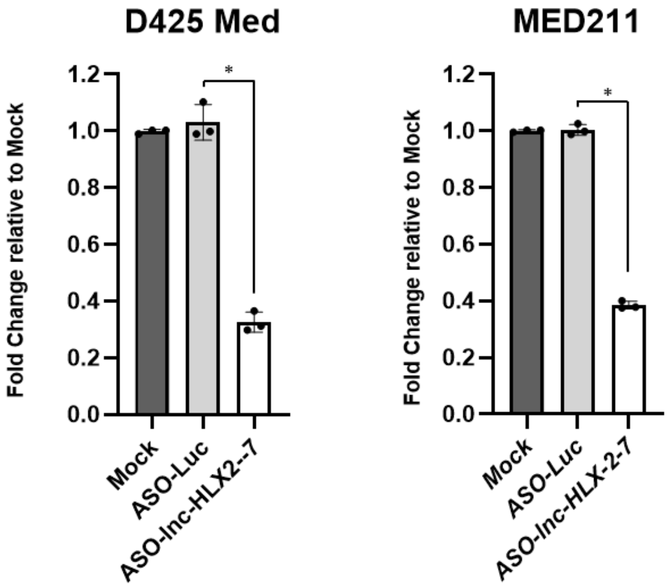


**Supplementary Figure 3. *lnc-HLX-2-7* regulates the expression of *HLX* coding gene.**

(A) Expression level of *HLX* in D425 Med and MED211 cells treated with ASO against the indicated genes in the x-axis. Relative expression level to mock is indicated in the y-axis. *$p$<0.01, Kruskal–Wallis analysis.

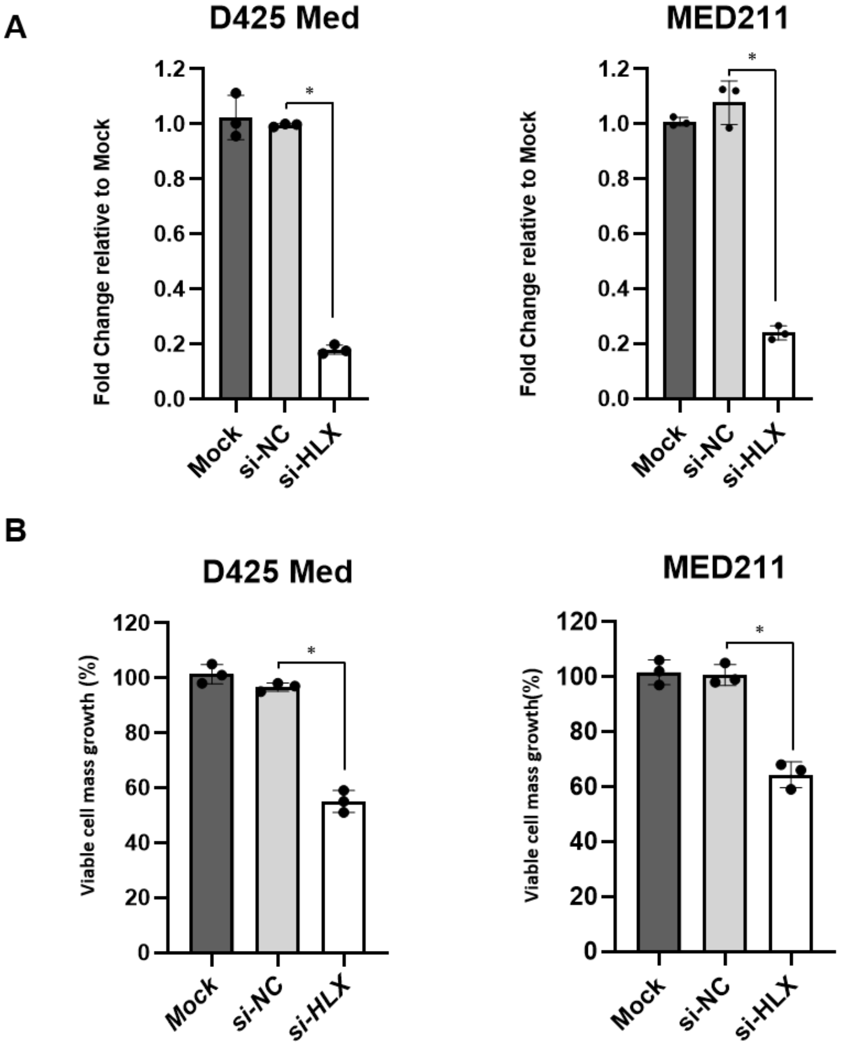


**Supplementary Figure 4. Effects of *HLX* expression on the proliferation of D425 Med and MED211.**

(A) Expression level of *HLX* in D425 Med and MED211 cells treated with siRNA against the indicated genes in the x-axis. (B) Viable cell numbers in D425 Med and MED211 cells treated with either si-*NC* or si-*HLX*. Relative value to mock is indicated in the y-axis. *$p$<0.01, Kruskal–Wallis analysis

A

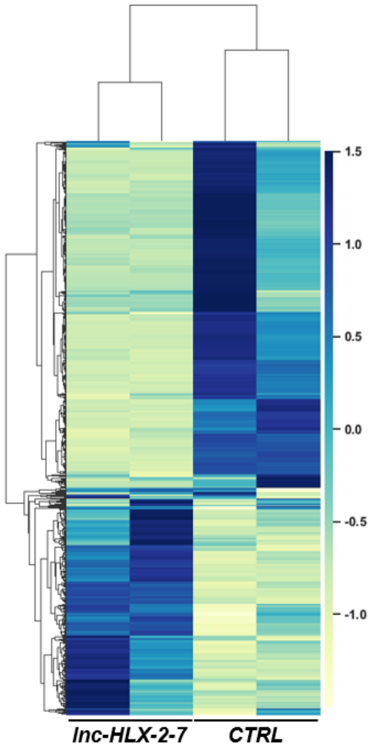


B

| Diseases or Functions Annotation | p-value | Predicted Activation State | Activation z-score |
|---|---|---|---|
| Cell death of osteosarcoma cells | 4.14E-08 | Increased | 3.500 |
| Cell death of cancer cells | 0.000348 | Increased | 2.807 |
| Cell death of tumor cells | 0.000425 | Increased | 2.338 |

C

| Upstream Regulator | MoleculeType | Predicted activation State | Activation z-score |
|---|---|---|---|
| MYCN | transcription regulator | Inhibited | -4.777 |
| MYC | transcription regulator | Inhibited | -4.448 |
| MLXIPL | transcription regulator | Inhibited | -4.264 |
| NFkB (complex) | complex | Inhibited | -2.889 |
| NFE2L2 | transcription regulator | Inhibited | -2.802 |
| 1,2-dithiol-3-thione | chemical reagent | Inhibited | -2.734 |
| KRAS | enzyme | Inhibited | -2.607 |
| HIF1A | transcription regulator | Inhibited | -2.531 |
| PKM | kinase | Inhibited | -2.425 |
| CD40LG | cytokine | Inhibited | -2.423 |
| ESR2 | ligand-dependent nuclear receptor | Inhibited | -2.414 |
| TNF | cytokine | Inhibited | -2.325 |
| EGFR | kinase | Inhibited | -2.093 |
| IL15 | cytokine | Inhibited | -2.030 |

**Supplementary Figure 5. *lnc-HLX2-7* interacting pathway genes in D425 Med cell.**

(A) Heatmap representation of genes up and downregulated after *lnc-HLX2-7* depletion in D425 Med cell ($p < 0.05$). (B) Molecular and cellular functions and diseases associated with these genes. (C) The most significant upstream regulators inhibited by depletion of *lnc-HLX-2-7*.

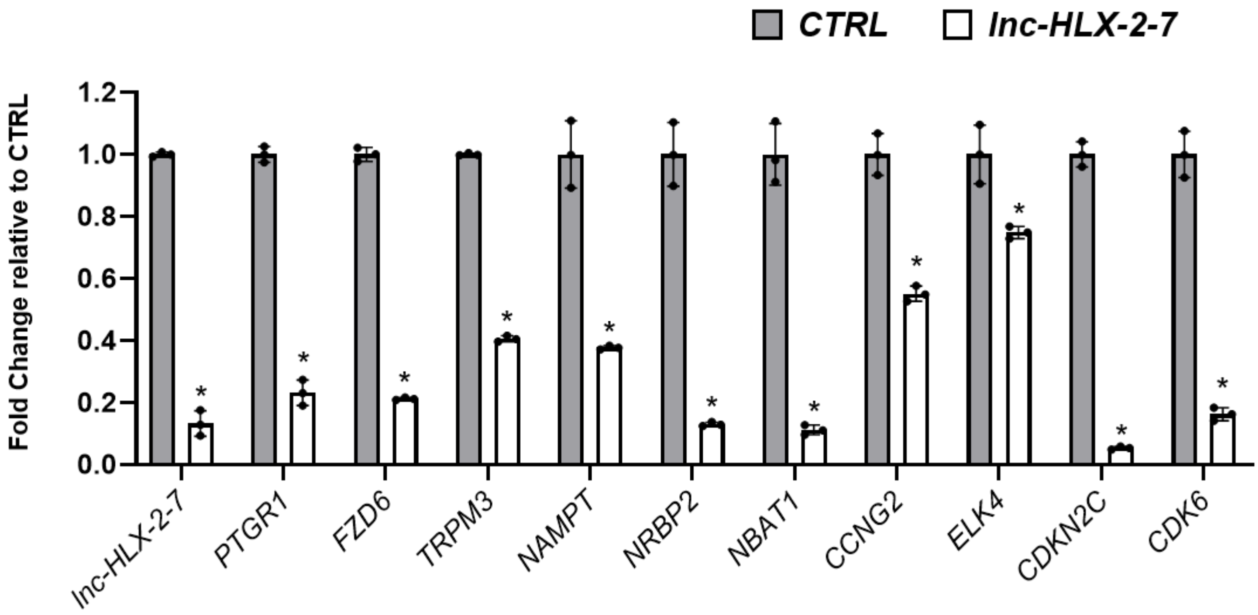


**Supplementary Figure 6. qPCR validation of D425 Med xenograft RNA-sequencing data.**

Expression levels of *lnc-HLX-2-7*, *PTGR1*, *FDZ6*, *TRPM*, *NAMPT*, *NRBP2*, *NBAT1*, *CCNG2*, *ELK4*, *CDKN2C* and *CDK6* were examined by qPCR in D425 Med xenograft. Relative expression levels compared with that in the CTRL tumor are indicated on the y-axis (n=3). Error bars indicate s.e.m. *$p < 0.01$, Student's *t*-test.

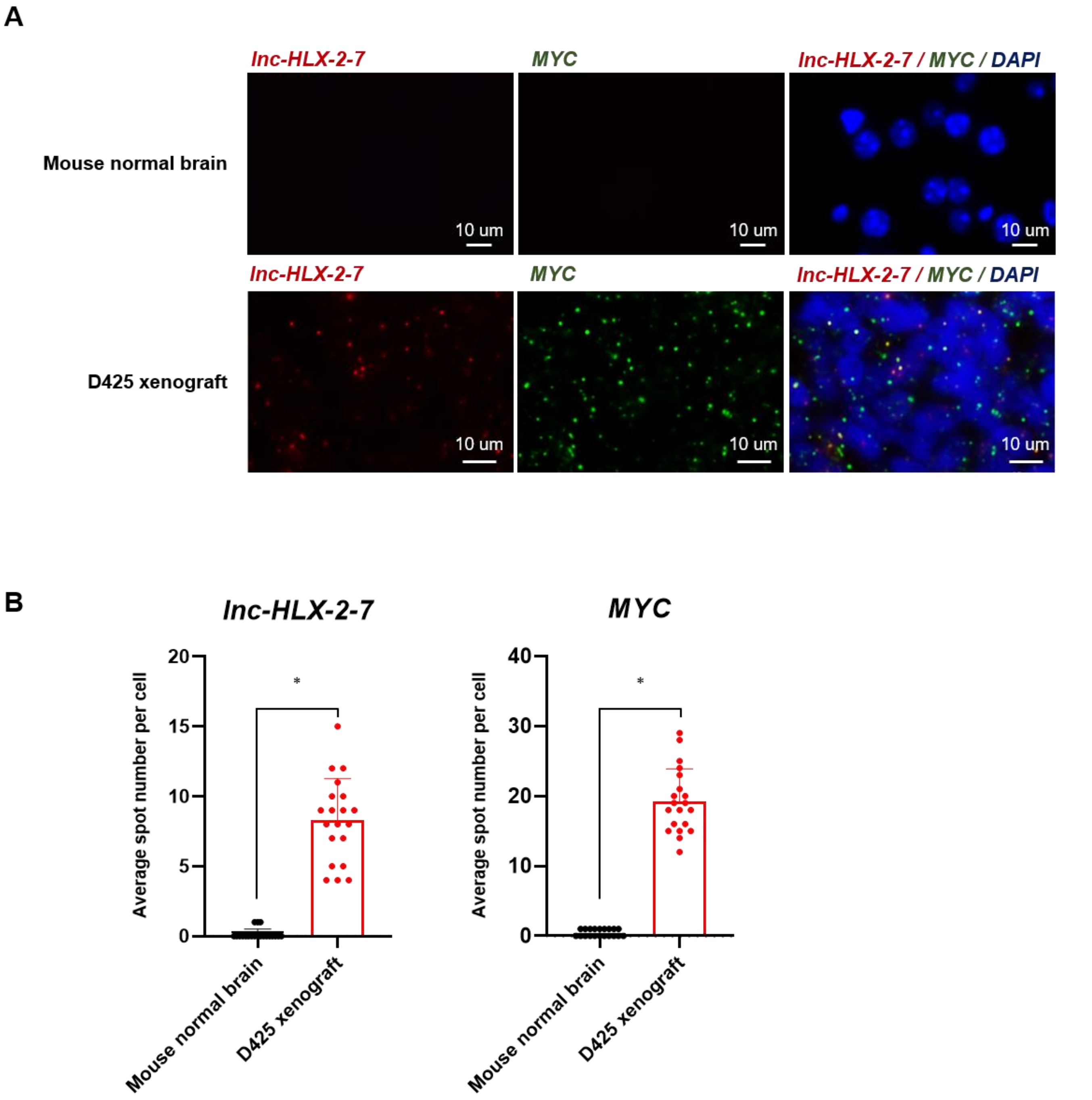
A
lnc-HLX-2-7
MYC
lnc-HLX-2-7 / MYC / DAPI
Mouse normal brain
D425 xenograft
10 um
B
lnc-HLX-2-7
MYC
Average spot number per cell
Mouse normal brain
D425 xenograft
*
0
5
10
15
20
30
40

**Supplementary Figure 7. Confirmation of the specificity of the *lnc-HLX-2-7* probe.**

(A) Representation of RNA-FISH analysis of *lnc-HLX-2-7* and *MYC* in MB tissues. RNA-FISH analysis of *lnc-HLX-2-7* and *MYC* in normal mouse brain (upper panels) and D425 xenograft (lower panels). Nuclei are stained with DAPI. Scale bars, 10 μm. (C) The spot numbers relating to *lnc-HLX-2-7* and *MYC* were quantified per cell in normal mouse brain and D425 xenograft. *$p$<0.01, Student's *t*-test.

**A**

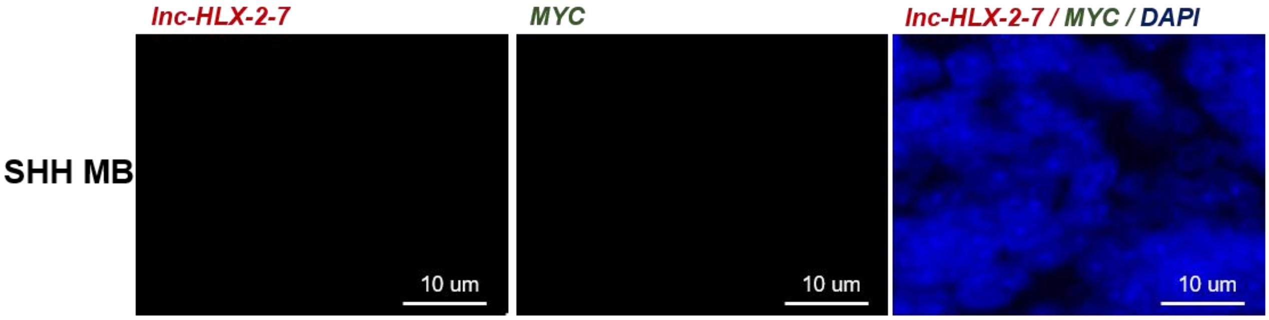
lnc-HLX-2-7
MYC
lnc-HLX-2-7 / MYC / DAPI
SHH MB
10 um
10 um
10 um

**B**

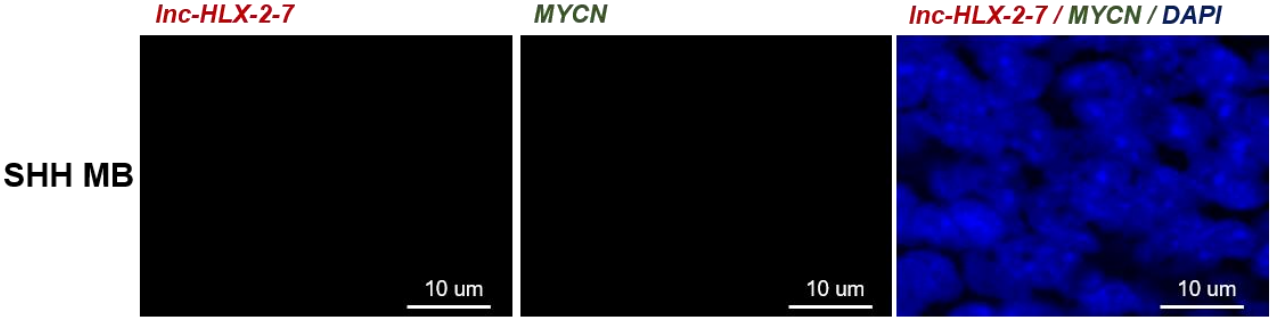
lnc-HLX-2-7
MYCN
lnc-HLX-2-7 / MYCN / DAPI
SHH MB
10 um
10 um
10 um

**C**

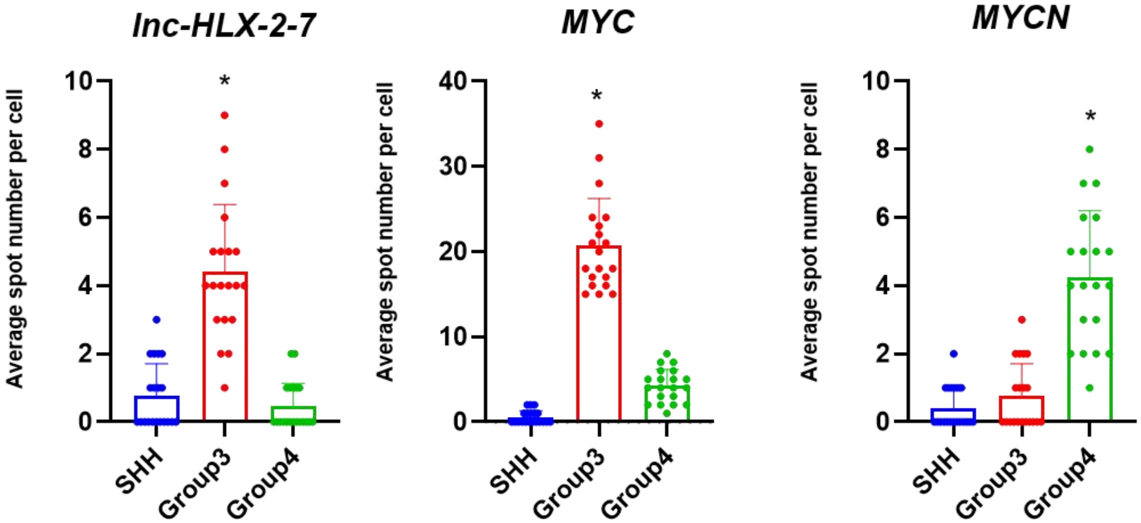
lnc-HLX-2-7
MYC
MYCN
Average spot number per cell
SHH
Group3
Group4
*

**Supplementary Figure 8. RNA-FISH confirms that *lnc-HLX-2-7* is not expressed in SHH MB patients.**

RNA-FISH analysis of *lnc-HLX-2-7* and *MYC* (A) or *MYCN* (B) in SHH MB tissues. Nuclei are stained with DAPI. Scale bars, 10 μm. (C) The spot numbers relating to *lnc-HLX-2-7*, *MYC* and *MYCN* were quantified per cell in Group 3, Group 4 and SHH MB patients. n=20, *$p<0.01$, Kruskal–Wallis analysis.
