## Supplementary methods for "The long non-coding RNA *lnc-HLX-2-7* is oncogenic in group 3 medulloblastomas"

### Isolation of single cells from orthotopic xenografts

30 days after of the injection of D425 Med cells and D425 Med cells with *lnc-HLX-2-7* deleted into the cerebellums, tumors were harvested and dissociated using a brain tumor dissociation kit (Miltenyi Biotech Inc., Auburn, CA) according to the manufacturer's protocol. To enrich human cells, mouse cells were depleted from the dissociated tumor cells using a mouse cell depletion kit (Miltenyi Biotech Inc.). The dissociated tumor cells were further sorted using a FACSAria (Beckton Dickinson, Franklin Lakes, NJ) to obtain live and singlet cells. The cells were resuspended in DPBS with 0.04% BSA to a final concentration of $1x10^6$ cells per ml.

### Processing of scRNA-seq data

Single-cell RNA-seq samples were classified into host and graft reads using XenoCell [1] and Xenome v1.0.1 [2]. The proportions of graft and host reads were 92.25% and 0.43% for D425, and 86.54% and 2.96% for *lnc-HLX-2-7* deleted D425 respectively. The remaining reads were classified as both, neither, or ambiguous. FASTQ files for graft were aligned to human genome hg38, indexed with GENCODE human annotations v34 [3] and augmented with lncRNA annotations from LNCipedia v5.2[4], using 10X Genomics *cellranger count* (https://support.10xgenomics.com/) and STAR v 2.7.0d_0221 [5]. For downstream integrated analysis, both samples were combined and normalized for the number of mapped reads per cell across libraries using 10X Genomics *cellranger aggr* function. 5,547 and 10,039 cells were detected for D425 and *lnc-HLX-2-7* deleted D425 respectively with post-normalization mean number of 18,034 reads per cell and median of 960 genes detected per cell.

**Quality control and clustering analysis of scRNA-seq data**

Quality control and clustering of scRNA-seq data were performed using Seurat v3.1.2[6] in R v3.6.1. Low quality and doublet cells were filtered by selecting cells with <10% mitochondrial percentage and expressing 200-2500 genes. 3,442 and 6,193 cells were retained for D425 Med and *lnc-HLX-2-7* deleted-xenograft samples after filtering. The count matrices for D425 and *lnc-HLX-2-7* deleted xenograft were normalized and integrated using *FindIntegrationAnchors* and *IntegrateData* functions. Principle component analysis (PCA) was subsequently performed. For combined clustering, 15 PCs with resolution=0.5 were used to obtain 5 clusters. The marker genes associated with each cluster were identified by finding differentially expressed features across clusters and using log2 fold change cutoff of ±0.2 and adjusted p-value of 0.05.
